# An Extreme Value Theory Model of Cross-Modal Sensory Information Integration in Modulation of Vertebrate Visual System Functions

**DOI:** 10.1101/423038

**Authors:** Sreya Banerjee, Walter J. Scheirer, Lei Li

**Affiliations:** Department of Computer Science and Engineering, University of Notre Dame, Notre Dame, Indiana, USA; Department of Biological Sciences, University of Notre Dame, Notre Dame, Indiana, USA

## Abstract

We propose a computational model of vision that describes the integration of cross-modal sensory information between the olfactory and visual systems in zebrafish based on the principles of the statistical extreme value theory. The integration of olfacto-retinal information is mediated by the centrifugal pathway that originates from the olfactory bulb and terminates in the neural retina. Motivation for using extreme value theory stems from physiological evidence suggesting that extremes and not the mean of the cell responses direct cellular activity in the vertebrate brain. We argue that the visual system, as measured by retinal ganglion cell responses in spikes/sec, follows an extreme value process for sensory integration and the increase in visual sensitivity from the olfactory input can be better modeled using extreme value distributions. As zebrafish maintains high evolutionary proximity to mammals, our model can be extended to other vertebrates as well.

## Introduction

The brain perceives the external world through an integration of stimuli received from different sensory modalities like vision, olfaction, and audition via the centrifugal pathway. A recent study taking inspiration from Cajal’s original work on brain mapping [18] describes current knowledge of the centrifugal olfactory and visual pathways in mammalian species as being incomplete. While, for instance, the signaling pathways mediating brain feedback in human olfaction have been characterized, the origins and effects of signals to visual system functions remain to be examined. In this work, we seek to understand the modulation of the circuits between sensory modalities. A crucial observation, yielding from our own work, points to how due to olfacto-visual sensory integration, measures of visual performance or behavior in response to multi-sensory input are enhanced, when a stimulus in one modality is ambiguous or undetermined. In fact, in all vertebrate species (e.g., teleost, reptiles, birds, rodents, primates) examined thus far, the retina receives brain feedback through the centrifugal visual pathways [17,19,33]. Depending on the species under consideration, the centrifugal pathways may originate from different parts of brain, such as the pre-tectal cortex, isthmo-optic nucleus, thalamus, or olfactory bulb.

In zebrafish *(Danio rerio*), the olfacto-retinal centrifugal (ORC) pathway originates from terminalis neurons (TNs) in the olfactory bulb (OB) and terminates in retina. TNs (Fig. 5A) synthesize gonadotropin-releasing hormone (GnRH) as a major neurotransmitter. In the retina, TN fibers synapse with dopaminergic interplexiform cells (DA-IPCs), retinal ganglion cells (RGCs), and possibly other retinal cell types. Insights from relatively recent research [22,28] have shown that the function of the ORC pathway is regulated by the olfactory input. TN input alters GnRH signaling transduction and decreases dopamine release in the retina, thereby increasing outer retinal sensitivity and inner retinal activity (e.g., firing of ganglion cells). Specifically, the olfactory input mediated by the ORC pathway decreases the light threshold (i.e., the minimum light intensity required to fire evoked action potentials) of retinal ganglion cells, and thereby increases retinal sensitivity. Together, the olfactory input amplifies behavioral visual sensitivity [31].

Zebrafish maintain high evolutionary proximity to mammals, and their retinas share great similarities to humans (e.g., structure, cellular organization, neural circuitry and signaling transmission) [27,50]. While much progress has been made to understand the anatomy of cross-modal circuitry in zebrafish, our knowledge of the underlying regulatory mechanism and physiological roles of centrifugal input to the retina is still in its nascent stage. Interestingly, Huang et al. [22] demonstrate how the visual sensitivity in zebrafish is increased in the presence of olfactory signals whereas disrupting the ORC pathway impairs visual function. An important observation found in that work reveals the importance of olfactory signals for vision. According to Huang et al., under normal conditions the minimum threshold light intensity to invoke a retinal ganglion cell response (measured in spikes/sec) in a dark-adapted zebrafish embryo may decrease 1-2 log units after olfactory stimulation. This demonstrates the sometimes dramatic impact of olfactory signals on vision.

Such a sudden gain in visual sensitivity through olfactory stimulation is an intriguing target for a computational model. We argue that visual sensitivity follows the statistical Extreme Value Theory (EVT). The mean visual sensitivity does not clearly explain the increased sensitivity due to olfactory signals since that scenario is able to sense a stimulus that is an extreme aberration from the norm, i.e., retinal ganglion cell responses without any olfactory stimulation. EVT lays solid groundwork for modeling as it is independent of the underlying distribution of data (all of the cell responses) and is only applicable to the tails of the distribution (the extremes) such that samples which have the least, or no possible, probability of occurrence under a central tendency model are distinguished, providing greater discrimination while requiring few statistical assumptions.

At a deeper level, one can ask the following question: is there a theoretical justification for using EVT for neural modeling? Our key insight is that the characterization of the firing behavior of a neuron as repeated integration/thresholding within a circuit suggests positive answers to these questions. Neurons are generally modeled as an electro-chemical process integrating input (ions) and eventually crossing a threshold whereby they “fire” and release ions. We posit that this inherently leads to an EVT-based model because the distribution of samples that exceed a threshold *T* likely yields an extreme value distribution (EVD). If all neurons use a fixed threshold *T*, the inputs to subsequent neurons in the circuit must follow an EVD, with each neuron integrating data from such a distribution and thresholding it. Thus, EVT can provide a plausible consistent multi-layer neuron model.

Beyond the merits of cultivating a better understanding of the operation of cross-modal sensory information integration in vertebrates, there is the possibility that an accurate computational model for this phenomenon could translate into a general algorithm for pattern recognition tasks in computer science. A direct application of this method lies in the development of novel information fusion algorithms that leverage inputs from multiple sensory modalities, i.e, vision and audition [34]. Another practical application is the invention of innovative sensors capable of detecting changes in the environment and then re-configuring on the fly to change operational parameters and power consumption requirements. Currently, sensors are typically designed to sense a single type of physical property such as temperature, pressure, radiation, motion or proximity. But with a biologically-consistent model they could be remodelled to use multiple observations from the environment for more agile operation. The work presented in this article is in this spirit of leveraging biological observations to forward engineer algorithms that can operate in a general context.

In the following sections, we provide a detailed explanation of our work. Section 2 describes the single unit cell recording procedure from which our analysis is derived and the definition of EVT from which the proposed model is based. Section 3 goes on to describe the exact specification of that model. Section 4 describes our experiments and Section 5 presents the corresponding results. Finally Section 6 concludes by putting this research into a larger biological and computational context.

## Materials and Methods

In this section, we explain the methods we use that are crucial for understanding our computational model of cross-modal sensory information integration. This includes the physical experiments that were conducted to collect the source data, as well as the formal elements of EVT.

### 0.1 Single-Unit Recordings and Odor Stimulation

This research builds upon the previous work of Huang et al. [22]. An overview is provided in Fig. 5B. Traces of RGC are recorded before and after odor stimulation (the sites of treatment are indicated by numbers 1 and 2 in panel B), or when dopamine and/or GnRH signaling transduction is manipulated by the application of receptor agonists or antagonists (indicated by numbers 3-8 in panel B). For electrophysiological recordings, zebrafish were anaesthetized with 0.04% 3-amino benzoic acid and immobilized by intraperitoneal injections of 3 - 5 *μ*l of 0.5 mg ml^−1^ gallamine triethiodide dissolved in phosphate-buffered saline (PBS), and then placed on a wet sponge with most of the body covered by a wet paper towel. A slow stream of system water (distilled water with ocean salt added, 3 g gal^−1^, pH 7.0) was directed into the mouth to keep the fish oxygenized. The eye was slightly pulled out of its socket and held in place by glass rods, thus exposing the optic nerve. Single-unit RGC responses (determined by the spike waveform) were recorded from the optic nerve by using a Tungsten microelectrode (resistance, 5-10 MΩ). Electrical signals were filtered with a band pass filter between 30 and 3000 Hz.

To test the effect of olfactory stimulation on visual sensitivity, we measured the light threshold required to evoke RGC responses before and after olfactory stimulation. Each fish was dark adapted for 30 min before the first RGC recording was made. The light stimuli (full-field dim white light, generated by a halogen bulb) were directed to the fish eye via a mirror system. The intensity of the unattenuated light beam (log *I* = 0) measured in front of the fish eye was 670 *μ*W cm^−2^ (Optical Power Meter, UDT Instruments, MD, USA). To determine the threshold, the light intensity was first set below threshold level (e.g., log *I* = −6.0) and then increased by 0.5 log-unit steps until the first light-evoked RGC responses were recorded (criteria, 20% above or below the rate of spontaneous firing). This light intensity was noted as the threshold. For each recording, 10 stimuli (600 ms flashes) were delivered at 3 s intervals.

Amino acids (methionine) were chosen to stimulate the olfactory neurons and thereby activate the ORC pathway. Previous studies have demonstrated that amino acids are strong odors for zebrafish [11]. Among the amino acids tested in zebrafish, methionine produced the most obvious and dose-dependent responses on visual function. In this study, odors (methionine, 0.5, 2, and 5 mM; total 8-10 *μ*l per stimulation) were delivered to the nostril through a glass pipette. The light threshold required to evoke RGC responses was measured before the application of methionine, and was measured again within 10 s following the application of methionine. Thereafter, the measurement was repeated at 1 min intervals for 10 min. In total, 24 cells were recorded. 24 animals were used in this process with 1 cell/animal for the recordings. Among these 24 animals, in response to odor stimulation, 17 showed increased visual sensitivity. In the remaining 7 animals, 6 showed no changes in visual sensitivity and 1 showed decreased visual sensitivity.

### Extreme Value Theory

The extreme value theorem [6] that underpins EVT (Fig. 5C) is very similar to the central limit theorem [24]. Both theorems involve limiting behaviors of distributions of independent and identically distributed random variables as *n*, the number of random variables, tends to ∞. However while the central limit theorem is concerned with the behavior of entire distributions of random variables, the extreme value theorem only applies to the random variables at the tails of those distributions.

To state this difference precisely, if *x_1_, x_2_*, …, *x_n_* represent the i.i.d. random variables from a distribution, then the central limit theorem describes the limiting behavior of *x_1_*, *x_2_*, …, *x_n_* while the extreme value theorem describes the limiting behavior of the extremes: max(*x_1_*,*x_2_*,…, *x_n_*) or min(*x_1_,x_2_*,…, *x_n_*) [6]. It encompasses a number of distributions that apply to extrema.

An extreme value distribution is a limiting model for the maximums and minimums of a dataset. A limiting distribution simply models how large (or small) the data to be modelled will probably get. It is widely used in applications where we are not only interested in estimating the average but also estimating the maximum or minimum [5,16,51,52]. For example, when designing a dam, engineers might not only be interested in the average yearly flood which foretells the amount of water to be stored in the reservoir, but also in the maximum flood, the maximum intensity of earthquakes in the region during the past decade, or maximum strength of concrete to be used in building the dam to avert the possibility of a disaster and for risk management. Castillo et al. [5] list a number of applications where extreme value distribution can be used.

Now that the preliminaries have been covered, we can formally define an extreme value theorem [13]:

*Let (s_1_, s_2_, …, s_n_) be a sequence of independent and identically distributed samples and let M_n_ = max(s_1_, s_2_, …, s_n_). If a sequence of pairs of real numbers (a_n_, b_n_) exists such that each a_n_ > 0 and*

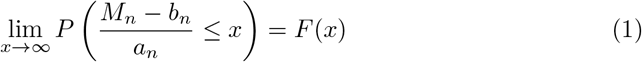

*then if F(x) is a non-degenerate distribution function, it belongs to one of three extreme value distributions: Gumbel, Frechet or Reverse Weibull*.

In contrast to the Gumbel or Frechet distributions which are used for unbounded data, the Weibull distribution applies to data that are bounded from below and when the shape (*k*) and scale (*λ*) parameters are positive (the Reverse Weibull is simply the opposite of the Weibull’s non-degenerate distribution function). Moreover, the Weibull is used for modeling minima. In order to use it for modeling data that fall in the upper tail of distribution, a minor adjustment needs to be made by flipping data based on the maximum value and then applying Weibull distribution. The probability distribution function of the two-parameter Weibull distribution is given as:

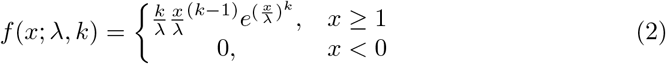

Note that there are other types of extreme value theorems one can make use of, such as the Pickands-Balkema-de Haam Theorem [38]. We limit ourselves to the theorem in Eq. 1 in this work for the modeling of explicit tail data, but we will invoke the Pareto distribution, which is derived from the Pickands-Balkema-de Haam Theorem, in the modeling of the overall distribution. This is described below in the next section.

## A Model for Cross-Modal Sensory Information Integration

Now that the relevant background has been introduced, we formally define our computational model for cross modal sensory information integration (Fig.2) It is motivated by the following hypothesis: *The tuning curves for RGC responses with and without olfactory signals are different. The extreme values lying near the tails of the distributions underlying those curves contribute to the determination of the visual sensitivity of zebrafish and should not be discarded as outliers.*

**Figure 1.**
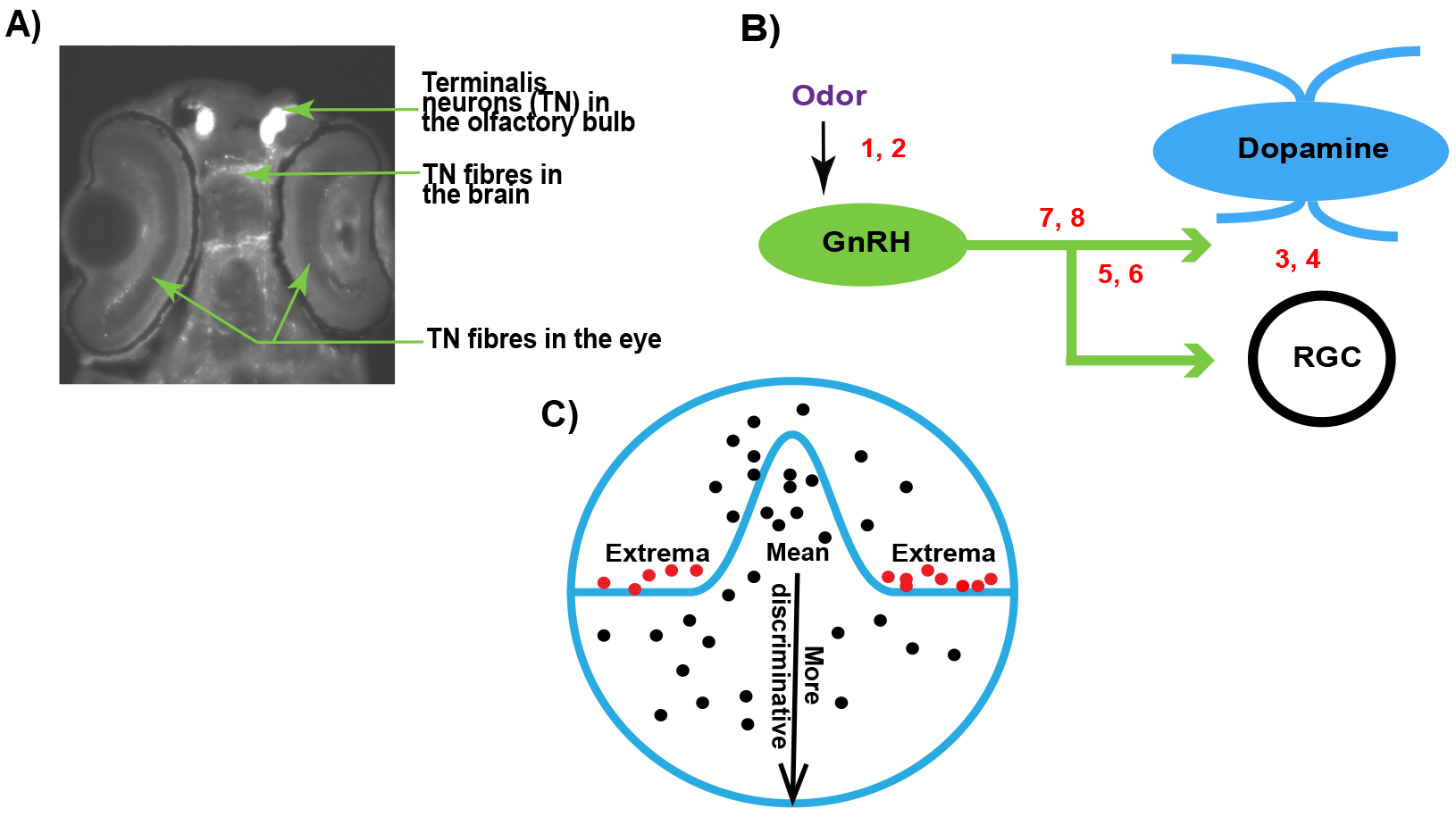
**(A)** A fluorescent image of a zebrafish (dorsal view, anterior is up) showing the terminalis neurons and axons **(B)** Schematic of the experimental setup for RGC recordings in response to olfactory and TN stimulation. The numbers correspond with the following conditions: 1,2 - sham or odor stimulation; 3,4 - activation or inhibition of dopamine receptors; 5,6 - activation or inhibition of GnRH receptors; 7,8 - manipulation of dopamine and GnRH receptors. **(C)** An overview of EVT. Prior work [15,26,49] suggests that it may be the extremes (red dots), and not the mean (black dots near the center of the circle), that produce strong responses in the brain.

**Figure 2.**
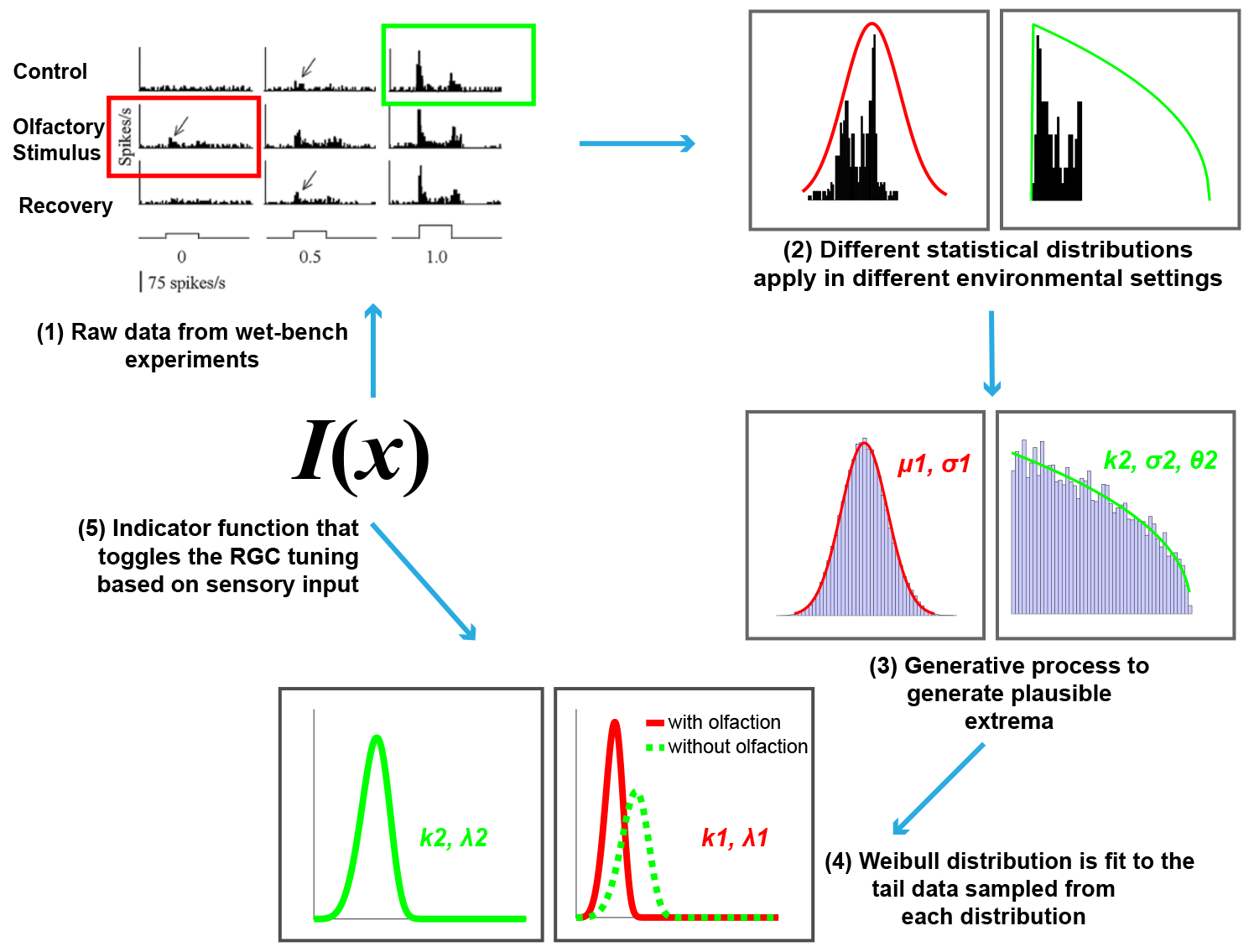
An overview of the proposed computational model of cross-modal sensory information integration. The red curve represents the RGC responses when an olfactory stimulus is present and the green curve represents the responses when there is no olfactory stimulus present. The first step represents the data collection effort from the wet-bench experiments. The second and third steps include fitting distributions to the data collected in order to draw samples for further processing. The fourth step represents fitting a Weibull distribution in order to model the underlying difference between visual sensitivity when olfactory input (i.e., additional sensory information) is present as opposed to when it is not. The final step is identifying an indicator function, *I(x)* that toggles between the two distributions based on the sensory input received.

The single unit recordings that we used for our experiments can be regarded as samples from a large population. One way to infer more about the population statistics is to extrapolate from the available samples by fitting distributions to them and sampling additional data. However, fitting a known distribution to available data can be difficult because of limited sample sizes, leaving one to make a “best guess” based on prior information about the behavior of large sample statistics. The best guess can come from making an assumption (for example, a null hypothesis as a starting place), or a more rigorous method of model selection using some metric.

If *n* represents the sample size, *n* ⟶ ∞ with the number of RGC responses acquired from an animal as it senses its environment over time. And the distribution of mean RGC responses calculated throughout an animal’s entire lifecycle becomes Gaussian. This assumption directly follows from the central limit theorem. So perhaps the underlying distribution of measured responses is also Gaussian (a typical assumption in such modeling). Because our experiments involve two different sets of RGC responses, with and without olfaction, we can hypothesize that each set is normally distributed with varying parameters. This null hypothesis can be tested through commonly used measures of normality, failing which it can be rejected and we can look for alternative distributions using a model selection approach.

In statistical modeling, statisticians are often faced with the task of selecting a suitable model (a distribution, in our case) among a set of viable and finite candidates. There are several metrics or selection criteria one can use to determine the best explanatory model given the data. The Bayesian Information Criterion (BIC) [35,43] serves as a canonical method for model selection when priors are hard to state precisely. In a large sample setting the model found by BIC is equivalent to the candidate model that is *a posteriori* most probable, given the available data. It primarily amounts to maximizing the likelihood function separately for each candidate model and then choosing the one for which the log likelihood is the largest, with a fixed penalty term for guessing the wrong model.

To identify a good distribution to fit to non-normally distributed empirical data, we used a Matlab implementation of BIC^1^. A large set of valid parametric distributions were fit to the data and sorted using the output of the BIC metric to compare the goodness of the fits. The overall process returns a set of fitted distributions with their respective parameters. The list of distributions that were tried includes: Beta, Birnbaum-Saunders, Exponential, Extreme Value, Gamma, Generalized Extreme Value, Generalized Pareto, Inverse Gaussian, Logistic, Log-Logistic, Log-Normal, Nakagami, Rayleigh, Rician, t Location-Scale, and Weibull. It was assumed that all data were continuous.

Our initial assumption that the overall data representing RGC responses without olfactory signals are normally distributed was rejected by the normality tests at the 1% significance level (a detailed description of the normality tests is given in Section 4). Using the BIC method, the distribution that fit accurately to the overall RGC response data without olfactory stimulation was found to be the Generalized Pareto distribution. Interestingly, this distribution is considered to be in the EVT family. The null hypothesis that the overall RGC responses with olfactory stimulus are normally distributed was not rejected at the 1% significance level by the normality tests, thus we fit a Gaussian distribution to that data.

Suppose we have *n* observations, or number of RGC responses. If *x_i_* represents the *i*-th RGC response where *i* ∈ (1, 2, …, *n*), the population statistics (mean *μ* and variance *σ*^2^) of the RGC response data with olfactory signal are found as the unbiased estimates of the distribution parameters and are given by the following equations:

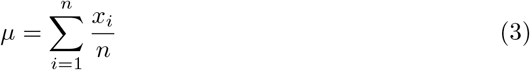

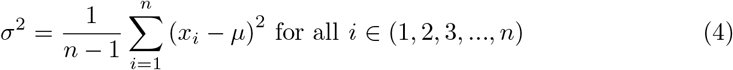

The probability density function for the Generalized Pareto distribution with shape parameter *k*, scale parameter *σ* and threshold parameter *τ* is given by the following equation:

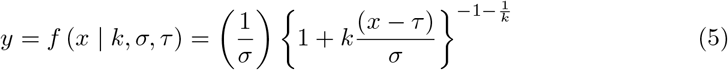

We used maximum likelihood to estimate the parameters *k* and *σ* from the two-parameter Generalized Pareto distribution by fitting RGC responses without olfaction^2^.

Having access to a model of the entire population facilitates generative sampling, which in turn allows for better tail modeling, and support for heightened visual sensitivity under certain conditions. Such generative processes in the brain may be responsible for a number of different phenomena, as they facilitate generalization in learning from limited sampling [39]. We use random sampling and the Metropolis-Hastings algorithm, a Markov chain Monte Carlo (MCMC) sampling method [20] to generate in total 100, 000 simulated RGC responses with and without olfaction respectively. The maximum (or the minimum) RGC response values within these samples follow an EVD. For our analysis, we concentrate only on the maximum RGC responses from the distributions described above because the lowest possible RGC response can be 0 spikes per second, indicating no response. Since the RGC responses (both with and without olfactory signals) can be assumed to be i.i.d samples from continuous distributions that are bounded from below, the Weibull distribution is the correct choice for modelling them. We expect the Weibull cumulative distribution curves (CDFs) for RGC responses with and without olfaction to be widely separated and the threshold RGC response value for an olfactory signal to shift sensitivity leftward (see Fig. 3 for an example), indicating that the cells are now more sensitive. This effect, replicated within the model, would confirm in a more rigorous sense that the presence of olfactory signals increases the fish’s sensitivity towards its surrounding and almost endows it with night vision that would be otherwise impossible in absence of those signals.

**Figure 3.**
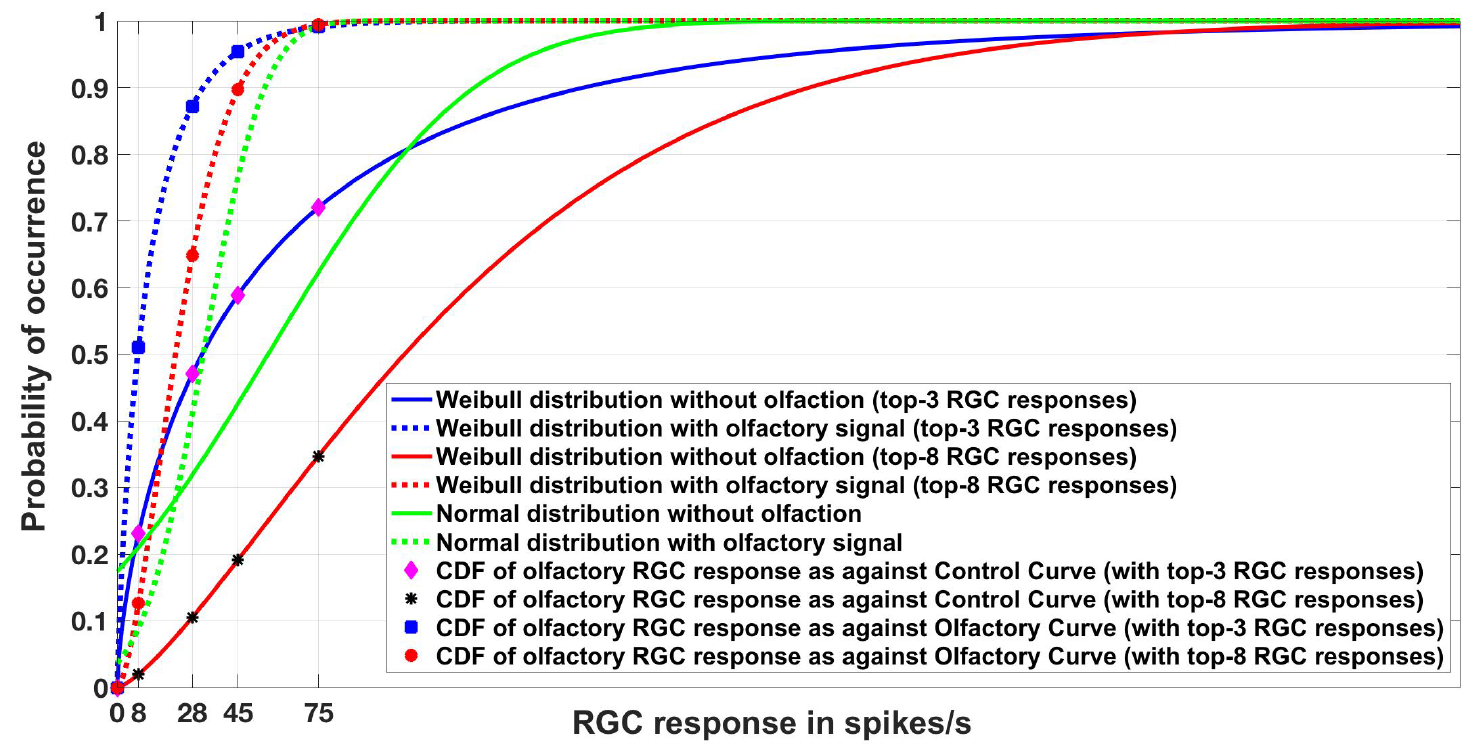
Experiment 1. Cumulative Distribution Functions for zebrafish with and without olfactory stimulation at light intensity 10^−^5 & 10^−^6 respectively. The curves depict the difference between central tendency modeling (green) and EVT modeling (red and blue). As can be seen, tuning becomes more sensitive when the Weibull distribution is used. The number of maximal RGC responses taken is either 3 or 8 (indicated within the parentheses). Best viewed in color.

This process is analogous to the super-additivity phenomenon in the multi-sensory superior colliculus of higher-order organisms like mammals, where the presence of two weak sensory signals from the environment enhances the animal’s neural response towards that environment [21]. The RGC threshold value represents an average RGC response for visual sensitivity, which changes throughout an animal’s entire life-cycle as it adapts to an ever-changing environment. However, the threshold varies (decreases or increases) in the presence or absence of a sensory stimulus other than visual input. This leads us to the possibility of the existence of some decision making mechanism in the fish’s brain that toggles between two different distributions to adjust the tuning of the RGCs based on sensory input. Mathematically, this decision making procedure can be implemented as an indicator function *I(x)*. If *θ* represents the parameters of an RGC distribution, i.e., the prior information available for RGC responses with or without olfactory signals and x represents a new RGC response due to a stimulus from the environment such that *x* ∈ *R^n^* (here *n* = 22, as we successfully retrieved 22 dimensions representing RGC spikes over time after stimulation of the olfactory neurons from the wet-bench experiments of Huang et al. For further explanation, see Section. 4), then the indicator function *I(x)* can be represented as:

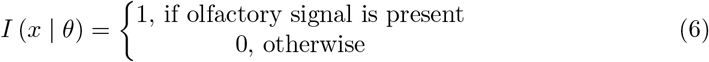

We speculate that the actual neural computation for the overall phenomenon is far more complex and is not restricted to just two modalities. However, given the recordings available for this study, we limit our model to just one particular circuit.

### Choices for an Indicator Function

For the indicator function, we address the following problem: given a set of vectors representing RGC responses in spikes/s with and without olfactory signals, is it possible for an indicator function to identify whether a new RGC response has been triggered after an olfactory signal or not? Our intuition behind using an indicator function is that such a process exists in some capacity in the brain where the presence of one signal enhances the other signal, thereby eliciting responses much different from the situation when the signal is not present. In essence, this task can be formulated as a binary classification problem with two possible outcomes: presence or absence of olfactory signals. Ideally, any discrimitative supervised learning method can easily solve the problem. For our analysis, we examine the utility of support vector machines and an artificial neural network which, to some extent, mimics the functions of a biological neuron and is closer to the mechanism that the brain uses to process such signals. The motivation for choosing these particular classifiers is their simplicity — we desire an indicator function with an efficient training regime that can operate over thousands of multi-dimensional data points, such as a large collection of RGC responses. Other classifiers (e.g., decision trees, random forests, logistic regression) may also be suitable.

#### Support Vector Machine

The Support Vector Machines (SVM) is a supervised learning approach that is widely used for classification and regression analysis. Since our data is numeric and high-dimensional, SVM [9] is a natural choice as it has been found to be extremely efficient in high-dimensional spaces for large-scale classification problems. SVMs use a subset of training points in the decision function, which form the “support vectors” that define the decision boundary between classes. As a consequence, it has been found to be memory efficient and has faster execution times if the data is normalized. For analysis, we assumed our data to be linearly separable and used a linear SVM kernel. We normalize our data using min-max normalization.

An SVM model with a set of labelled training data tries to find an optimal hyperplane for classifying new samples based on some constraints. Given a training dataset, *D* = (*x_i_, y_i_*) of size *m* with *x_i_* = (*x_1_, x_2_*, …, *x_n_*), an n-dimensional feature/attribute vector and label, *y_i_* = −1 or +1, formally the SVM classifier can be defined as a quadratic optimization problem solving the following equation:

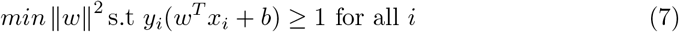

where *w* = (*w_1_, w_2_*, …, *w_n_*) is a weight vector and *b* is the bias.

An important consideration when designing a SVM model is the parameter *C* that dictates the trade-off between having a wide margin and correctly classifying training data.

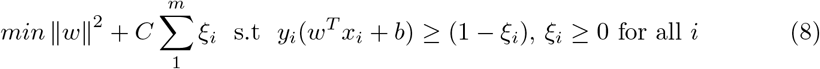

A larger value of *C* implies a smaller number of mis-classified training samples and is prone to overfitting.

#### Artificial Neural Network

We also consider a multi layer perceptron (MLP) neural network as the indicator function. Similar to SVM, MLP is a supervised learning algorithm that learns a non-linear mapping from input, *x* ∈ *R^n^* where *n* represents the number of dimensions, to *y* ∈ *R^m^* where *m* can be any number *m* < *n*, depending on the number of classes in the training dataset. However, unlike SVMs, a simple MLP includes one or more hidden layers consisting of artificial neurons. The hidden layers act as feature detectors and gradually discover the salient features of the training data through backpropagation [40, 53]. Each neuron includes a non-linear and differential activation function and is connected to every neuron in the previous layer exhibiting a high degree of connectivity between layers. As a result, due to the distributed nature of non-linearities, the learning process is difficult to visualize. However, neural networks are usually assumed to be non-parametric functions, i.e., they can be used as function approximators without having any prior information about the distribution of input or training dataset and hence are well suited to represent the indicator function. If *x* represents a *p*-dimensional input vector such that *x* = (*x_1_*,*x_2_*,*x_3_*, …, *x_p_*) with *y* = (+1, —1) as labels and *g* : *R* ⟶ *R* as the activation function, then the equation of a single neuron is given by:

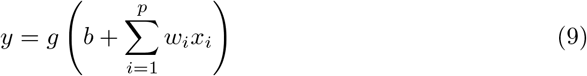

where *w* = [*w_1_, w_2_, w_3_*…*w_p_*] represents the weights learned through backpropagation.

## Experiments

### Data collection and representation

As stated above, the first step in building a computational model of this nature is to attempt to define the underlying distribution of the data one is trying to explore. We use the data from a study by Huang et al. [22] for our analysis. The data consists of single unit RGC responses measured in spikes/sec before and after olfactory stimulation under varying light intensity (see Fig. 2 from Huang et al.). In terms of raw data organization, it is primarily a histogram with the x-axis representing the visual sensitivity of fish binned into approximately 22 positions representing a timestamp and their corresponding frequency measured in spikes/sec on the y-axis. Under normal conditions, the minimum threshold light intensity to invoke a retinal ganglion cell response in a dark-adapted zebrafish embryo is 10^−5^. However, with olfactory stimulation with methionine, the threshold light intensity decreases to 10^−6^. We calculated the minimum RGC response threshold to be at 75 spikes/sec. Hence, the data can be separated into two parts: one with olfactory stimulus and the other without it. In total, there were 22 RGC responses across time with olfactory stimulus and 29 without olfactory stimulation.

### Experiment 1

The first experiment was to check whether the raw data we collected from the experiments confirms our hypothesis that the EVT can be applied to build an accurate model. We posit that since the RGC responses with olfactory stimulation represent extreme aberration from the baseline and are non-negative integers, the Weibull distribution is the right candidate for modeling our data. But how differently does our data fit with the Weibull distribution versus a central tendency model like the Gaussian distribution? We explore this by comparing the CDFs of the Weibull and Gaussian distributions with parameters derived from our data.

### Tests of normality and synthetic data generation

Using the data collected from wet-bench experiments as a basis, we simulated an expansive data space by fitting distributions over the original data. The goal was to generate as much evidence as possible for statistical inference. However, in order to fit distributions to generate more samples from the existing data, we need to make some assumptions about the underlying distribution. Initially, as described above in Section 3, we assumed a null hypothesis that the distribution of RGC responses in a zebrafish throughout its entire lifecycle is Gaussian. Since our work involves two different sets of RGC responses — one with olfactory stimulus and the other without it — under this assumption the distributions underlying each should be Gaussian with different parameters. To test this, we performed several commonly used tests of normality: the Kolmogorov Smirnov test [32], the Shapiro-Wilk test [44], and a Lilliefors test [7,29,30]^3^. Due to the small sample size (*n* = 22 or 29), we preferred the Shapiro-Wilk test over Kolmogorov-Smirnov and Lilliefors. For datasets that failed the normality test, The BIC selection criterion was deployed to find another distribution with the best fit. Afterwards, we generated 100, 000 non-negative samples of RGC responses from the respective distributions for further analysis.

### Experiment 2

The second experiment was to check whether the points we sampled confirm our hypothesis that the EVT can be applied in a generative scenario. In order to verify this, we fit a Weibull distribution to the top *n* RGC responses to understand how the the curves vary when olfactory input is present as opposed to when it is not. The value *n* was selected via empirical observation. The sampling methods used were: random sampling and MCMC sampling. Since EVDs like the Weibull only apply to samples at the tails of distributions, it is independent of the underlying distribution of the data as a whole. Hence, irrespective of the overall data distribution and sampling process, the results of Experiment 2 for the Weibull distributions for the top *n* responses should ideally be similar to Experiment 1. We expect the Weibull cumulative distribution functions for data with and without olfactory stimulus to be widely separated, with the curve for data with olfaction shifting leftward, giving higher probability scores to RGC responses that would be improbable under conditions where olfaction is not engaged.

### Experiment 3

Additionally, we wanted to corroborate whether we can define a deterministic indicator function such that given some RGC response it is possible for the function to identify if an olfactory stimulus is present or not. In essence, this task becomes a binary classification problem where the presence of olfactory signals can be labelled as 1 and the absence as 0. As described above in Section 3, we use a linear SVM or a multi-layer perceptron as our binary classifier. For consistency in the operation of the indicator function, we limit the dimensionality of all vectors to the dimensionality of RGC responses with olfactory stimulus (*n* = 22). We use the 100,000 samples we generated for each scenario (with olfactory stimulus and without olfactory stimulus), dividing the sets into 80% training and 20% testing partitions.

In summary, the entire modeling effort is encapsulated in the following steps (also depicted in Fig. 1):

1. **Data collection and representation.** This step consists of collecting and representing data based on the wet-bench experiments for control (without any stimulation) and experimental (with olfactory stimulation) zebrafish as a histogram and collecting the statistics for further analysis.
2. **Experiment 1.** This first test consists of an experiment to evaluate our hypothesis that EVT applies with the raw data collected in step 1. We fit Gaussian distributions (to the entire collection of data with and without olfaction individually) and Weibull distributions (to the top-*n* RGC responses from the two datasets). The value *n* was selected via empirical observation.
3. **Tests of normality and synthetic data generation.** Here we begin by assuming that the distribution of RGC responses in a zebrafish throughout its entire life cycle is normal, and attempt to falsify that assumption via tests of normality. The appropriate distributions are subsequently fit to the data to generate 100, 000 synthetic samples. The data with olfactory stimulus follows the Gaussian distribution, whereas the underlying distribution for data without olfactory stimulus is Generalized Pareto.
4. **Experiment 2.** Similar to Experiment 1 but instead uses 100, 000 generated samples and only Weibull distributions fit to the top-*n* samples generated to examine how olfactory signals influence visual sensitivity as reflected by the CDF curves for the two scenarios. The value *n* is selected through empirical observation.
5. **Experiment 3.** This experiment involves identifying an indicator function *I(x)* that can distinguish when an olfactory stimulus is present and when it is not. Here this function is a deterministic binary classifier, either a linear SVM or a multi-layer perceptron.

## Results

### Experiment 1

Fig.depicts the result of Experiment 1 which was conducted to examine the difference between central tendency modeling and EVT modeling. The data for this experiment were what was directly collected from the wet-bench experiments for both control (without olfaction) and experimental (with olfaction) zebrafish.

As can be seen in the figure, with olfactory stimulation the visual sensitivity in zebrafish shifts leftward, making the RGC responses below the normal threshold of 75 spikes/s probable, as indicated by the physiological experiments of Huang et al. [22]. Moreover, if we look closely, the Weibull distributions (represented by the red and blue solid and dashed lines) are a better fit to the data because the RGC responses with olfactory stimulation represent the outliers or extreme responses as opposed to RGC responses without any stimulation. If we fix our attention at the threshold RGC response at 75 spikes/s, the Weibull curves provide a better explanation for getting an RGC response below 75 spikes/sec for olfactory stimulation in comparison to the normal distribution which almost makes those values improbable. In other words, the tuning becomes more sensitive if we use the Weibull distribution. We plotted the curves by varying *n* (*n* = 3, 8) of the top-*n* RGC responses. The tuning becomes more sensitive as *n* becomes smaller.

### Tests on normality and synthetic data generation

The null hypothesis that the data without olfactory stimulus are normally distributed was rejected at the 1% significance level for all of the tests. However, the other assumption of normality for data with olfactory stimulus was not rejected at the 1% significance level. Based on these results, we fit a Gaussian distribution to the data with olfactory stimulus. Using the BIC selection criterion to find the best fit, the distribution for the data without olfactory stimulus was determined to be Generalized Pareto. We then collected non-negative samples simulating RGC responses via random sampling or MCMC sampling (100, 000 samples from each sampling method), to be used for fitting a Weibull distribution to the top *n* samples in order to understand how the curves vary when olfactory input is present (i.e., when the overall distribution is Gaussian) as opposed to when it is not (i.e., when the overall distribution is Pareto).

### Experiment 2

Figs. 4 and 5 show the models of visual sensitivity calculated over the simulated data from random sampling and MCMC sampling^4^. Similar results are achieved for both sampling methods. An important observation to note here is that tuning is always more sensitive when olfactory stimulus is present. The values of *n* in this experiment are much larger (*n* = 50, 250) due to the increased availability of data, but still represent a small number of points from the tail of the overall distribution. The CDF curves for data with and without olfactory stimulation are widely separated and the width of separation increases as *n* grows larger. This reflects how the visual sensitivity threshold can change throughout a fish’s life cycle as it is exposed to an ever-changing environment and acquires new RGC responses for modulating its internalized model of visual sensitivity. Note that zebrafish build new cells within their nervous systems via a neurogenesis process, meaning the number of responses available at a point in time can change in a non-stimulus dependent way. Our proposed model supports this phenomenon.

**Figure 4.**
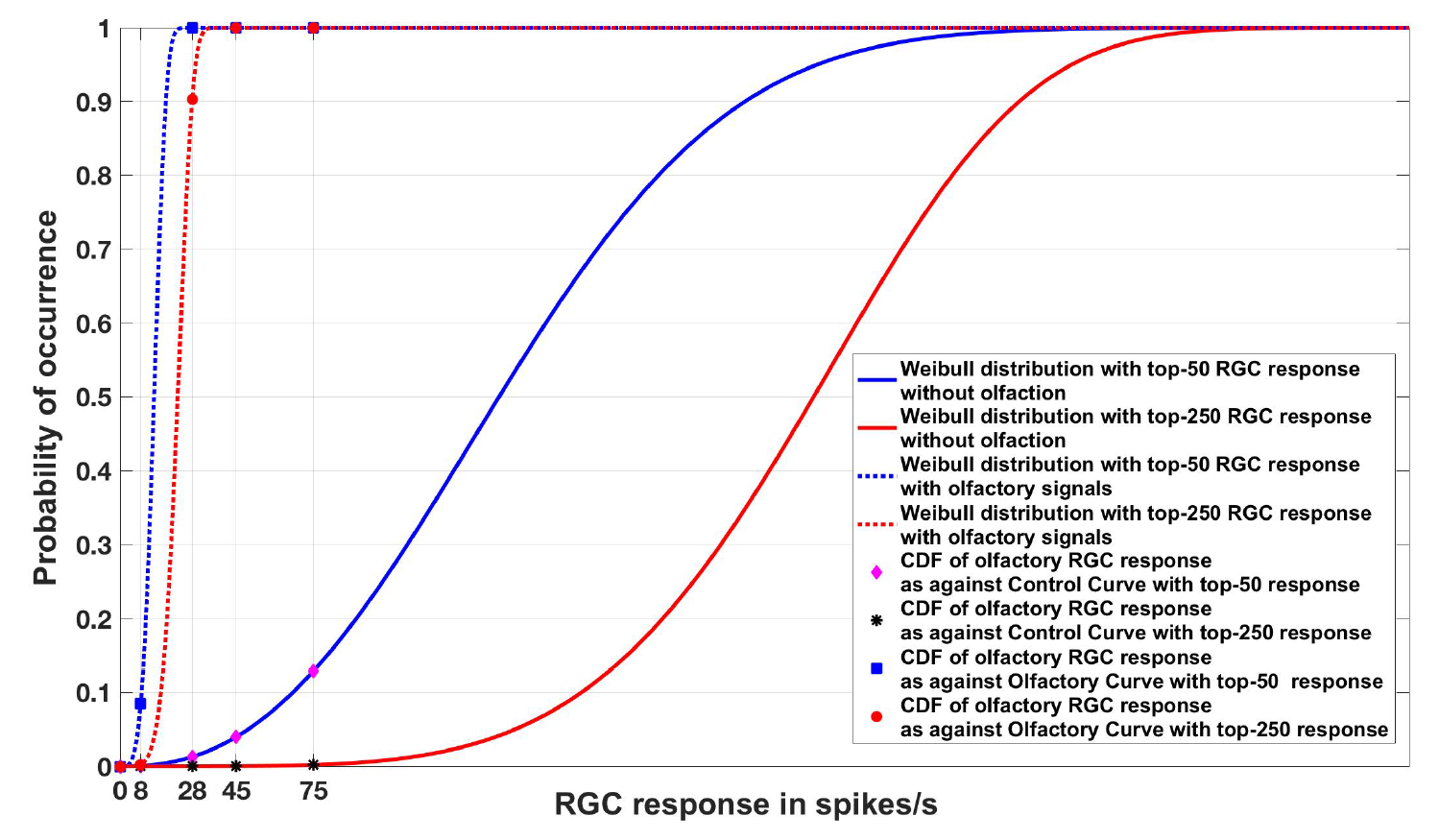
Experiment 2. Cumulative Distribution Functions for zebrafish with and without olfactory stimulation at light intensity 10^−5^ and 10^−6^ respectively, with data points generated through random sampling. The curves labeled “Control” in the legend describe the Weibull distributions (as represented by the solid blue and red lines) without olfactory stimulus. As can be seen, tuning is most sensitive when an olfactory stimulus is involved. Best viewed in color.

**Figure 5.**
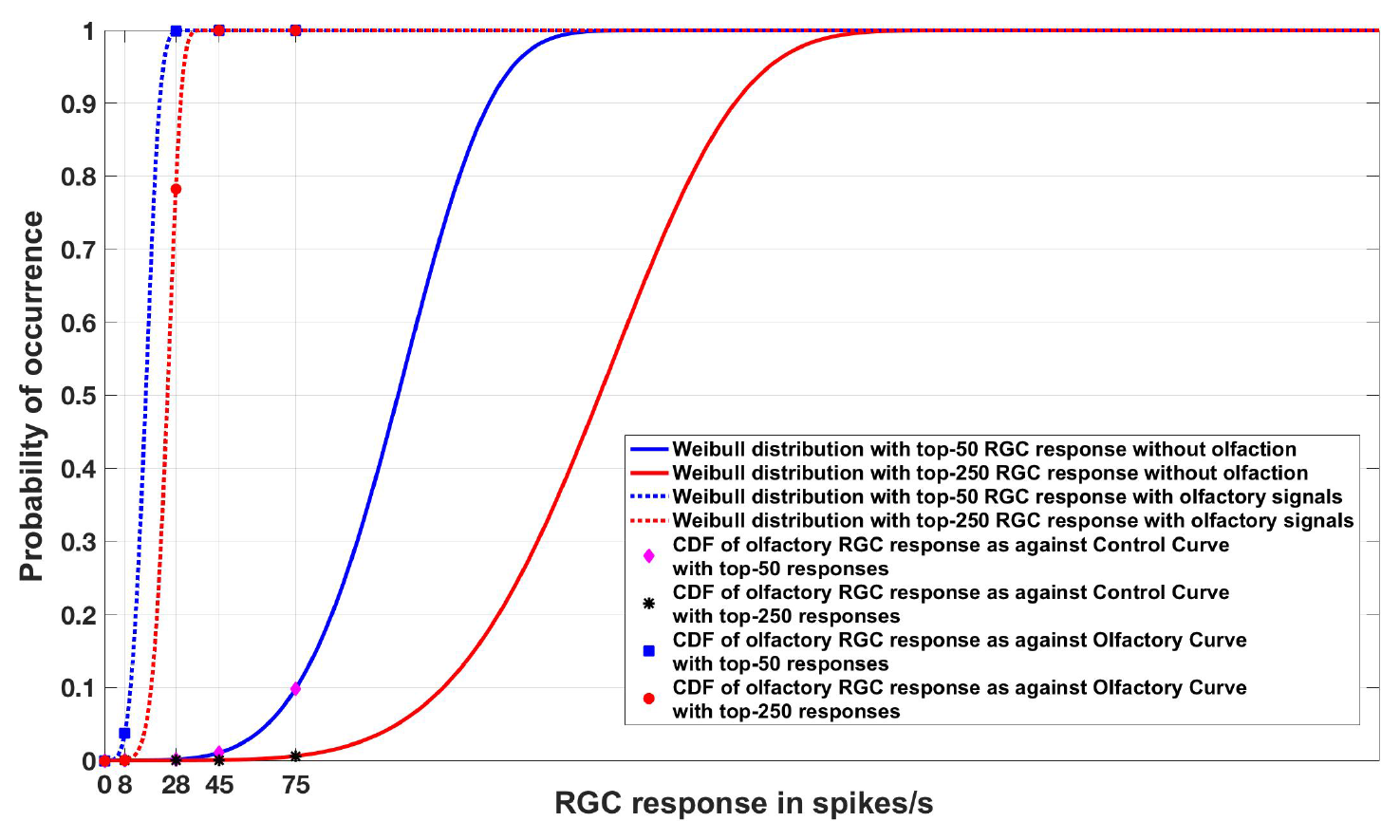
Experiment 2. Cumulative Distribution Functions for zebrafish with and without olfactory stimulation at light intensity 10^−5^ and 10^−6^ respectively with data points generated through MCMC sampling. The curves labeled “Control” in the legend describe the Weibull distributions (as represented by solid blue and red lines) without olfactory stimulus. The result is very similar to random sampling — the tuning is more sensitive when an olfactory stimulus is involved. Best viewed in color.

### Experiment 3

With respect to testing the possible indicator functions *I(x)*, we began by considering a linear binary SVM classifier trained using 80, 000 generated samples and tested using 20, 000 generated samples. With random sampling, we achieved a testing accuracy of 95.5 (+/- 0.163) percent, but with MCMC sampling accuracy decreased to 93.925 (+/- 0.123) percent. With a multi-layer perceptron classifier, the accuracy dropped to 95.25 (+/- 0.007) percent using the same training-testing split and data from MCMC sampling^5^. The success of this experiment establishes that the two different classes of RGC responses are separable. Thus it is possible, in a statistical learning sense, to have a mechanism to toggle between RGC tuning configurations when an olfactory stimulus is present and when it is not. One possibility for why the classification was successful in these experiments is that the indicator function implicitly learns that the data are distributed differently in the two classes (Generalized Pareto for data without olfactory stimulus and Gaussian for the data with olfactory stimulus). That the two classes of data are distributed differently lends further support to our hypothesis that an indicator function is involved in the integration of cross-modal sensory information — the distributional difference facilitates a very straightforward pattern recognition process to separate the classes.

## Discussion

As vertebrates evolved over centuries, sensory organs adapted with the ever-changing environment. In many vertebrate species, at any given time the brain integrates and processes multi-sensory information. In humans, for example, the functions of the olfactory and visual systems are influenced by sensory input from each organ. Most mammals have specialized multimodal neurons in the superior colliculus that are capable of integrating multiple stimuli from the environment and providing a uniform reaction. In lower vertebrates such as fish, however, such advanced mechanisms are absent. In zebrafish, the integration of sensory information from the olfactory system facilitates signaling transduction in the visual pathway. As a consequence, retinal neural activities such as the firing of retinal ganglion cells are increased. This is particularly important for wild type animals that live under natural environmental conditions. For example, zebrafish normally mate in the early morning hours before the sun comes up, during which time the light illumination is low. It is conceivable that under such conditions stimulation of olfactory neurons may increase visual sensitivity and thereby facilitate the process of mating. While the cellular mechanisms underlying this olfacto-retinal sensory integration have been well characterized, statistical models that describe the phenomenon at the cellular level have not been described. In this paper, we have described a computational model that supports the research into how the visual system integrates information from other sensory modalities.

While the cellular mechanisms underlying this olfacto-retinal sensory integration have been well characterized, statistical models that describe the phenomenon at the cellular level have not been described. In this paper, we describe a computational model that supports the research into how the visual system integrates information from other sensory modalities.

The idea of building computational models for multisensory input has been explored previously [1,2,10]. When it comes to determining the statistical relationship between sensory responses among different sensory organs, the Bayesian model has been a preferred framework. However, almost all of the existing work focuses on higher vertebrates such as mammals. Angelaki et al. [2] attempted to reconcile the difference between the traditional physiological studies on multisensory integration with computational and psychological studies using Bayesian inference on the visual-vestibular system for the perception of self-motion in macaques. They describe how the multimodal neurons represent probablistic information defined by multiple stimuli and propose that the special neurons accomplish near optimal cue integration through a linear summation of input signals.

With respect to models of simpler animals, Wessnitzer et al. [54] explore multimodal sensory integration for navigation from the physiological perspective of the insect’s nervous system. In zebrafish, using a similar linear model [23] the contribution of different types of cone photoreceptor cells to photopic spectral visual sensitivity was determined. This was done by re-modeling the electroretinographic data recorded from the cornea, which include absorbance spectrum of four types of cone photoreceptor cells (cone cells that are sensitive to ultra-violet light, blue light, green light, and red light, respectively) given as the visual pigment template for the appropriate maximum absorption, neural signals obtained from different cone cell types, relative fraction of the individual cone cells across the retina, and linear gains for each cone type [4]. The model incorporated the first-order cellular and biophysical aspects of cone photoreceptor cells and thereby predicts the second-order physiological functions of cone cell-mediated visual sensitivity. Using this model, linear gains that represent the strength of four different types of cone cell-derived neural signals onto four different inferred cone processes in the whole retina can be assessed.

Turning to extreme value theory, the objective of nearly all extant models in computational neuroscience has been to discard the extreme values located at the tails of distributions as noise and concentrate on the mean or average. However, evidence suggests that extremes, and not means, of cell responses direct activity in the brain. For example, the ability of primates, like macaque monkeys, to identify individual faces can be localized to a group of special neurons that fire in response to specific regions of face [15]. An interesting finding that came out of that study was that neurons were tuned to the geometry of extreme facial features. Previous investigations along this line (and a more recent one) concentrated on how the brain fundamentally adapts itself to the statistics of the sensory world, extracting relevant information from sensory inputs by modeling the distribution of inputs that are encountered by the organism [46, 47]. This lead to the advent of “sparse coding” which attempts to explain how neurons encode sensory information using a small number of active neurons at any given point in time [36]. A direct extension of this work suggests that sparse coding is an all-pervasive phenomenon used by all types of sensory neurons in different modalities across different species [37]. EVT builds upon these concepts but is more specialized.

Much prior work related to EVT modeling has focused on various non-biological applications from trend detection in ground-level ozone [48] to quantifying extreme precipitation levels using generalized Pareto distributions [8]. Other applications of EVT include, but are not limited to, finance, telecommunications and the environment [12], and hydrology [25]. Recent work in computer vision and machine learning has extensively used the concept of EVT [3, 14, 41, 42, 45]. For instance, for biometric verification systems, Shi et al. [45] used the General Pareto Distribution to model the genuine and impostor scores and made a significant observation that the tails of each score distribution contain the most relevant information that helps in defining each distribution considered for prediction and the associated decision boundaries, which are often difficult to model.

Our research extends this theory to multi-sensory inputs through a model that demonstrates strong neural fidelity. With a biologically-consistent information fusion algorithm based on retinal circuits in the zebrafish, we believe that we have access to a better general solution to the problem at hand and many other information processing problems of interest. Here, we develop a neural computation model that simulates the process of multi-organ sensory integration and predicts the consequence of sensory integration in higher-order brain functions. In contrast to Gaussian modeling, we propose that EVT models of the extrema found in the tails of the data can form a powerful basis for cross-modal sensory information integration, facilitating heightened sensitivity in targeted modalities that have been influenced by a stimulus in the environment. This resulted in the development of a computational EVT-based framework for multi-organ sensory integration in the zebrafish that is not only an explanatory model in neuroscience, but also shows promise for applications in machine learning and neuromorphic systems.

## Acknowledgments

The research was funded by the Department of Defense (Army Research Laboratory) under the contract, W911NF-18-1-0292.

## Author Contributions

WJS and LL initially conceived of the idea. LL was responsible for conducting the wet-bench experiments and preparing the source data. SB designed, analyzed, implemented the model and wrote the paper. WJS supervised the entire modelling effort.

## Data Availability Statement

The datasets analyzed for this study and the codes are released for reproducability and can be found in the [Zebrafish_EVT] [https://github.com/sbanerj2/Zebrafish_EVT].

## Ethics Approval Statement

An ethical review process was not required for our study. All data used in this article come from a previously published study [22]. All experimental procedures in that paper adhered to the NIH guidelines for animals in research.

## Footnotes

1 https://github.com/dcherian/tools/blob/master/misc/allfitdist.m

2 For finding the maximum likelihood estimates of the Generalized Pareto distribution, we used the Matlab function *gpfit*, which only returns the estimates of the shape *k* and scale *σ* parameters of a two-parameter Generalized Pareto distribution. The function *makedist* was then used to create a probability distribution object reflecting where samples are taken from, using the parameters *k* and *σ*.

3 We used the following Matlab implementations of the normality tests: *lillietest* (for the Lilliefors test), *swtest* (from Matlab central for the Shapiro-Wilk test), *kstest* (for the one-sample Kolmogorov-Smirnov test). Each of these tests returns a decision (1 or 0) for the null hypothesis that the data comes from a distribution in the normal family, against the alternative that it does not come from such a distribution. A result of 1 rejects the null hypothesis at the 5% significance level (default). For our experiments, we set the significance level to 1%.

4 We ran experiments 1 and 2 ten times. In each of those trials, the leftward shift of the distribution after olfactory stimulation was preserved.

5 Each of these experiments was run ten times. The numbers in parentheses represent standard error.

